# Disruption of the AgMOBP1–PfSyn5 interface abolishes malaria transmission

**DOI:** 10.64898/2026.05.01.722187

**Authors:** Julian Ramelow, Guodong Niu, Luka Kostallas, Liliana Lai, Xiaohong Wang, Jun Li

## Abstract

Malaria control remains constrained by an incomplete understanding of the molecular interactions that enable *Plasmodium falciparum* transmission within mosquito vectors. Although blood-stage parasite biology has been extensively studied, the mosquito-stage processes required for transmission are less well defined. Here, we characterize a previously unannotated *Anopheles gambiae* midgut factor, Midgut Ookinete Binding Protein 1 (AgMOBP1; AGAP008138), and define its role in parasite infection. AgMOBP1 is a membrane-associated protein localized predominantly to the apical surface of midgut epithelial cells, is induced by blood feeding, and appears restricted to anopheline mosquitoes. Functional infection assays showed that exogenous recombinant AgMOBP1 enhanced *Plasmodium* infectivity by increasing midgut oocyst burden, whereas anti-AgMOBP1 antibodies significantly reduced infection, establishing AgMOBP1 as a host factor that promotes transmission. Using biochemical interaction assays, we identified the parasite binding partner of AgMOBP1 as the *P. falciparum* syntaxin-5–like protein PfSyn5. Disruption of this interaction with anti-PfSyn5 polyclonal antibodies produced exceptionally potent transmission-blocking activity, with complete inhibition at 1 µg/mL and an estimated IC50 of ∼3 ng/mL (∼20 pM). This potency is approximately one million-fold lower than the total IgG concentration in normal human plasma. Together, these findings identify a critical mosquito–parasite molecular interface required for malaria transmission and establish the AgMOBP1–PfSyn5 axis as a promising target for transmission-blocking vaccines, monoclonal antibodies, and other intervention strategies.

**Significance Statement:** Interrupting malaria transmission requires a deeper mechanistic understanding of how *Plasmodium* parasites interact with their mosquito vectors. Here, we define a previously uncharacterized mosquito midgut factor, AgMOBP1, and identify its parasite binding partner, the *Plasmodium falciparum* membrane protein PfSyn5. We show that this molecular interaction is required for efficient parasite transmission to *Anopheles* mosquitoes. Remarkably, antibodies against PfSyn5 completely blocked transmission at low concentrations, revealing exceptional functional potency. These findings uncover a previously unknown mosquito–parasite invasion pathway and establish the AgMOBP1–PfSyn5 interface as a high-value target for transmission-blocking vaccines, monoclonal antibodies, and next-generation malaria control strategies.

## Introduction

Malaria remains one of the most consequential infectious diseases worldwide. Transmitted by *Anopheles* mosquitoes, malaria causes hundreds of millions of clinical cases annually and continues to impose substantial mortality and economic burden (*1*). According to the World Health Organization, an estimated 247 million malaria cases and 619,000 deaths occurred globally in 2022 (*1*), with ∼95% of cases and deaths concentrated in the African region. Among the human-infective *Plasmodium* species, *P.* f*alciparum* accounts for most severe disease and malaria-associated mortality, particularly in sub-Saharan Africa (*2, 3*).

Despite sustained global control efforts, malaria elimination remains elusive. The effectiveness of frontline interventions is increasingly threatened by the spread of antimalarial drug resistance in parasites and insecticide resistance in mosquito vectors (*4, 5*). These converging challenges underscore the need for complementary strategies that target parasite transmission in addition to disease treatment and prevention.

The *Plasmodium* life cycle includes an obligate developmental phase within the mosquito vector that is essential for onward transmission (*6, 7*). Following ingestion of infected blood, gametocytes rapidly differentiate into gametes within the mosquito midgut, where fertilization gives rise to mobile ookinetes. These ookinetes traverse the peritrophic matrix and midgut epithelium (*8*) before differentiating into oocysts beneath the basal lamina. Oocysts subsequently produce thousands of sporozoites that migrate through the hemolymph to the salivary glands and transmitted during a subsequent blood meal. Because only small fraction of parasites successfully complete development in the mosquito, this stage steps represents a major bottleneck and an attractive point for intervention (*9*).

Vaccination remains a central pillar of malaria control. Most vaccine efforts have focused on pre-erythrocytic or blood-stage parasites. The most advanced licensed malaria vaccine is RTS,S, targets the circumsporozoite surface protein (PfCSP) but provides only partial and time-limited protection (*10–12*). By contrast, transmission-blocking vaccines (TBVs) are designed to prevent parasite development within the mosquito and thereby reduce community-wide spread. Although TBVs do not directly protect vaccinated individuals from disease, they are increasingly recognized as critical components of eradication strategies. Progress in TBV development, however, has been limited by the small number of validated targets from mosquitoes and parasites and by the technical challenges of studying sexual-stage parasites (*13*).

The mosquito midgut is a highly specialized epithelial barrier (*14*) that plays a central role in parasite establishment. Following blood feeding, a glycan-rich peritrophic matrix (PM) forms around the blood bolus and separates ingested microbes from the epithelium (*15*). To establish infection, ookinetes must breach this barrier and interact directly with midgut cells (*16, 17*). Multiple, not mutually exclusive models have been proposed for ookinete traversal of the midgut, including intercellular (*18, 19*), paracellular (*18, 20*), and transient intracellular routes (*21*). Generally, the penetration of the parasite into the midgut wall occurs asynchronously, a period ranging from approximately 18 to 36 hours after blood-feeding (*22, 23*). Regardless of the precision route, successful invasion requires molecular interactions between parasite proteins and mosquito midgut factors.

Ookinete motility and invasion depend on specialized secretory organelles. Unlike other invasive stages, ookinetes lack rhoptries but are enriched in micronemes (*24, 25*), which release proteins required for adhesion, motility, and host cell interaction (*26, 27*). Several parasite proteins, including CTRP (*28*), CelTos (*29*), MAOP(*30*), and α-tubulin-1(*17*), contribute to mosquito infection. However, the specific receptor-ligand interactions that mediate parasite engagement with the mosquito midgut remain poorly defined.

Previous studies have identified mosquito proteins that either restrict or facilitate malaria transmission. While factors such as APL1 (*31*) and FBN30(*32*) inhibit parasite infection, others including AnAPN (*33, 34*), annexin protein (*35*), FREP1 (*16, 17*), and AgPfs47Rec (*36, 37*) promote transmission through distinct mechanisms. Expanding the repertoire of mosquito-stage transmission factors is therefore essential for the development of next-generation interventions.

Here, we characterize AgMOBP1, a previously uncharacterized mosquito protein that promotes malaria transmission (*38*). We show that AgMOBP1 is a membrane-associated factor enriched at the apical surface of midgut epithelial cells and induced by blood feeding. We further identify the parasite syntaxin-5–like protein PfSyn5 as its binding partner and demonstrate that disruption of the AgMOBP1–PfSyn5 interaction strongly impairs parasite transmission. Together, these findings define a previously unrecognized molecular axis governing mosquito-stage malaria infection and establish a promising target for transmission-blocking intervention.

## Results

### AgMOBP1 gene architecture, protein structure, expression profile, and contribution to *Plasmodium* infection

*AgMOBP1* gene is located on chromosome 3R of the *Anopheles gambiae* PEST genome (AgamP4_3R:6,092,088-6,093,806, reverse strand), contains three exons encoding a 510-amino-acid protein with a predicted molecular mass of 54.7kD and an isoelectric point of 4.44 (**Fig. 1A**). Sequence analysis identified N-terminal signal peptide (residues 1-22), consistent with entry into the secretory pathway, but no recognizable functional domains. Interpro predicted a large non-cytoplasmic region, whereas classical transmembrane helices and GPI-anchor motifs were absent. Notably, AgMOBP1 contains CVVA at position 479, a CaaX motif near the C-terminus for the typical post-translational modification site of a lipid-anchored membrane protein (*39*), suggesting possible prenylation-mediated membrane association.

**Figure 1.**
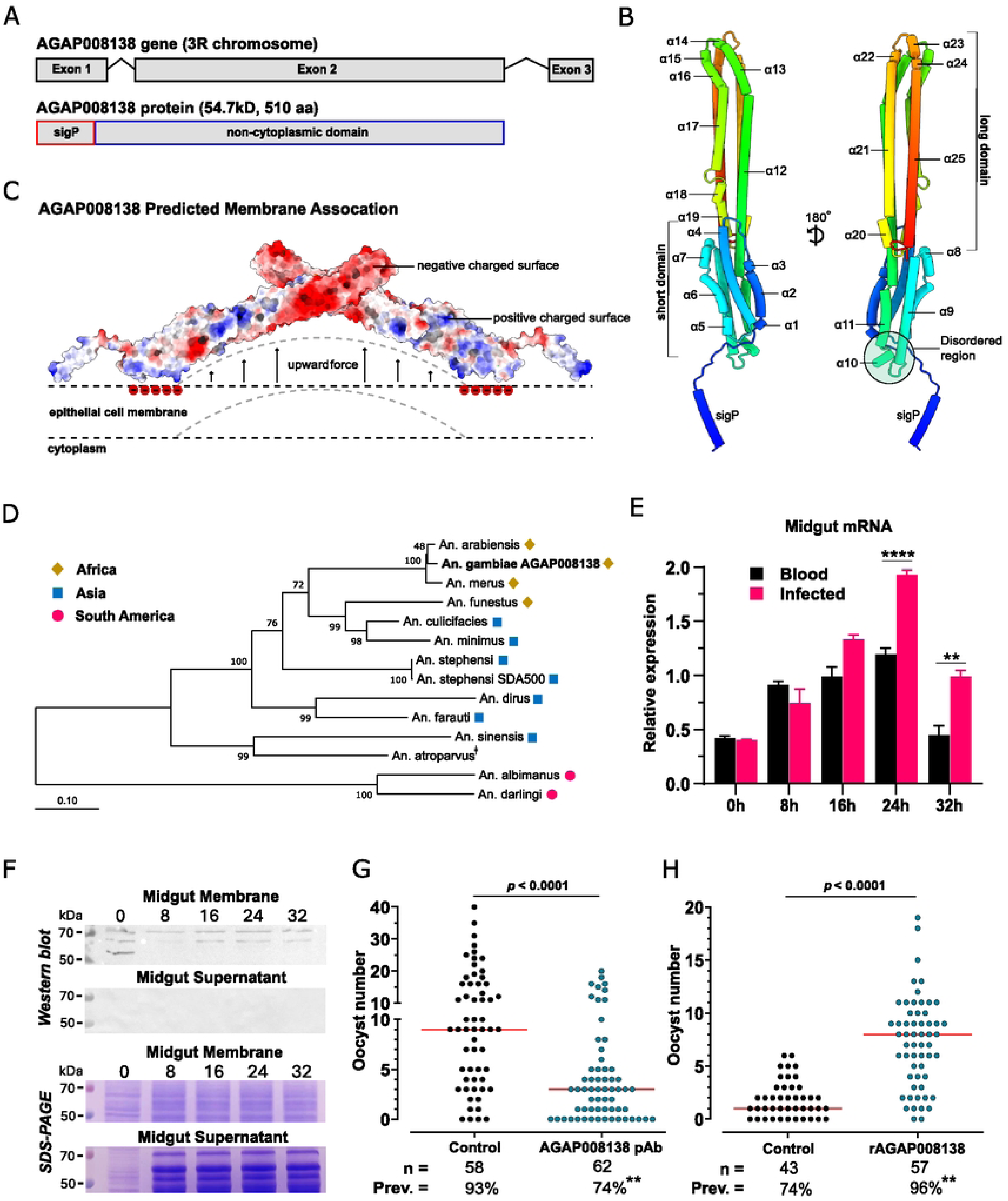
An overview of the gene structure, expression, and the influence of AgMOBP1 on *Plasmodium* infection. **A.)** AgMOBP1 consists of three exons on the 3R chromosome, creating a protein containing 510 amino acids with a signal peptide (sigP) with a predicted size of 54.7kD. **B.)** AlphaFold2 prediction of AgMOBP1 with an average of over 90% accuracy in rainbow coloring. AgMOBP1 has a short and a long domain consisting of 25 individual alpha-helices connected by coiled coils. A disordered region connects the short and long domains. **C.)** Predicted membrane association of AgMOBP1, anchored by lipids at the position of 479 on helix 25. Positive charges on the surface could also attract the negatively charged membrane heads. Combined with another AgMOBP1 protein, it creates an upward force that bends the membrane, creating an opening for vesicle fusion, which presents one of many possibilities. **D.)** Phylogenetic tree of AGAP008183 and its orthologs in other anophelines. There is a clear evolution pattern from anophelines present in Africa (gold diamonds) to Asia (blue square) and South America (pink circle). Bootstrap values higher than 45% (1,000 replications) are shown. The scale indicates the number of amino acid substitutions per site. **E.)** Relative expression of AgMOBP1 in *An. gambiae* midguts of blood-fed and *P. falciparum*-infected blood-fed mosquitoes at various time points was assessed by RT-PCR using the 40S S7 ribosomal protein gene as the internal reference. 0 h is the sugar-fed control baseline. **F.)** Western blot and SDS-PAGE of AgMOBP1 protein expression in *An. gambiae* midgut membrane (Upper) and midgut cytosol (midgut supernatant, Lower) from blood-fed mosquitoes 8, 16, 24, 32 h PBF, or sugar-fed (0 h). **G.)** *P. falciparum* oocyst infection levels in midguts from mosquitoes treated with anti-AgMOBP1 polyclonal antibodies (10 μg/ml) or control rabbit pAb (10 μg/ml). This represents one of three experiments that show similar results. **H.)** *P. falciparum* oocyst infection levels in midguts from mosquitoes treated with rAgMOBP1 protein (∼10 μg/ml on Hi5 cells) or control rGFP (∼10 μg/ml on Hi5 cells). *n*, number of mosquitoes analyzed, Prev., the prevalence of infection. \*\**p* < 0.01, χ^2^-squared test. We repeated the experiments three times, and the conclusions were the same.

Structural modeling predicted that AgMOBP1 is dominated by 25 α-helices and coiled-coil elements organized into two major domains (long and short) (**Fig. 1B**). Surface-charge analysis suggested asymmetric electrostatic properties across the molecule, and disorder prediction identified flexible internal segments (**Fig. 1C**). Together, these features are consistent with a scaffolding or membrane-interactive protein capable of mediating protein–protein or protein–membrane interactions rather than enzymatic catalysis (**Supplemental Fig. S1**).

Comparative sequence analyses further indicated that AgMOBP1 is evolutionarily restricted to anopheline mosquitoes. Orthologs were detected across multiple *Anopheles* species and followed expected phylogenetic relationships (**Fig. 1D)**. Sequence conservation was highest within the *An. gambiae* complex, intermediate in Asian vectors such as *An. stephensi*, and lower in New World species such as *An. albimanus*, consistent with lineage-specific divergence after speciation.

To determine whether AgMOBP1 expression is regulated by feeding or infection, we quantified transcript abundance in mosquito midguts. AgMOBP1 was expressed at low basal levels in sugar-fed females but was strongly induced after blood feeding, reaching peak expression at 24 h post-blood meal before returning toward baseline. A similar temporal pattern was observed after infectious blood meals; however, transcript levels were significantly higher in *P. falciparum*–infected blood-fed midguts than in uninfected blood-fed controls at 24 h and 32 h post-feeding (**Fig. 1E**).

Polyclonal antibodies raised against recombinant AgMOBP1 detected a major ∼55-kDa species and additional higher-molecular-weight bands in mosquito midgut extracts. Following blood feeding, the abundance of these bands changed dynamically, with the ∼65-kDa species closely paralleling the mRNA induction profile. Biochemical fractionation showed that AgMOBP1 was recovered in the membrane pellet but not in the soluble cytosolic fraction, supporting the prediction that AgMOBP1 is membrane associated despite lacking a canonical transmembrane domain (**Fig. 1F**).

We next tested whether AgMOBP1 functionally influences parasite infection. In standard membrane feeding assays, addition of anti-AgMOBP1 antibodies significantly reduced both oocyst intensity and infection prevalence relative to control antibodies (**Fig. 1G**) and this inhibition was concentration dependent (**Supplemental Fig. S2**). Conversely, supplementation with purified recombinant AgMOBP1 significantly increased infection burden at the highest concentration tested (**Fig. 1H**) and the promotion is concentration-dependent (**Supplemental Fig. S3**). Anti-AgMOBP1 antibodies did not cross-react with parasites (**Supplemental Fig. S4)**, indicating that the inhibitory effect was mediated through blockade of the mosquito factor rather than direct parasite neutralization. Collectively, these findings establish AgMOBP1 as a blood meal–induced mosquito midgut factor that promotes *Plasmodium* infection.

### Subcellular localization of AgMOBP1 in insect cells and in mosquito midguts

To define the subcellular distribution of AgMOBP1, we performed immunofluorescence assays (IFA) in baculovirus-infected insect cells and mosquito midguts (**Fig. 2**). In heterologous insect cells, recombinant AgMOBP1 localized predominantly to the cytoplasmic membrane, with prominent punctate and peak-like signal enrichment at the apical surface (**Fig. 2A**). Three-dimensional reconstruction confirmed a brush-like surface pattern (**Fig. 2B**), whereas control cells lacking AgMOBP1 expression showed no signal (**Supplemental Fig. S5**).

**Figure 2.**
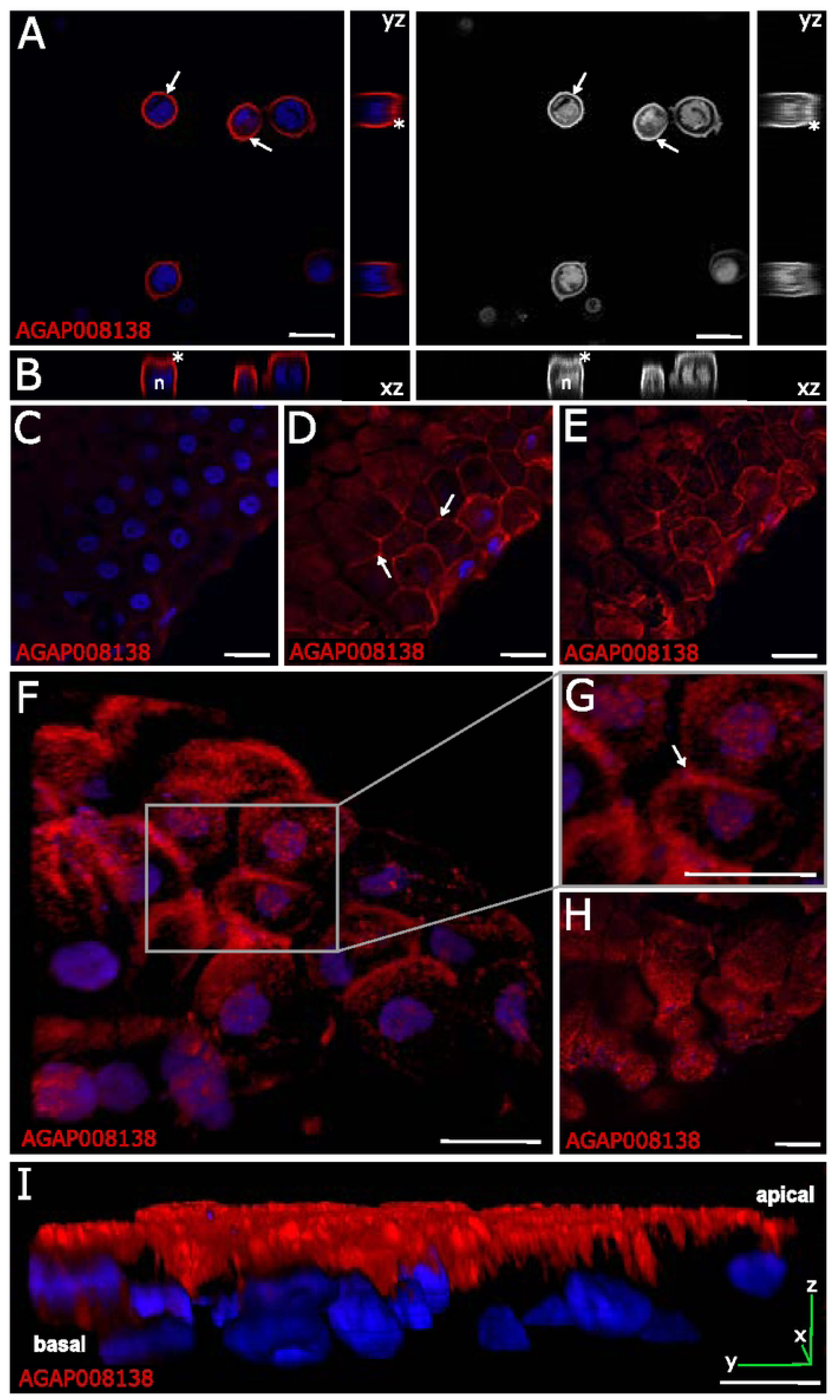
Immunofluorescence assays and confocal revealing the subcellular location of AgMOBP1 in recombinantly infected Hi-5 insect cells and *An. gambiae* midgut cells (24-28 h post-feeding with blood) to be in the cell membrane. The red/grey color depicts AgMOBP1 protein locations. **A.)** mid-cell section of Hi-5 cells with recombinant baculovirus infection, orthogonal views with yz (side) plane views, **B.)** 3D reconstruction and orthogonal views with xz (bottom) plane views, supporting AgMOBP1 protein on cytoplasmic membrane. * Denotes apical and top brush-like appearance. n, nucleus. AgMOBP1 (red) localization in blood-fed mosquitoes: **C,D,E.)** Midgut section at basal end, middle, and apical section, respectively, supporting AgMOBP1 proteins at midgut apical end, **F.)** 3D reconstruction looking at the apical side, **G.)** magnified apical side. **H.)** Another 3D view of the midgut looking at the apical side. **I.)** 3D side view with XYZ plane. White arrows indicate the presence and accumulation of AgMOBP1 at apical sites of a midgut. Scale bars, 20µm.

We also analyzed the tissue distribution of AgMOBP1 in mosquito midguts using IFA under a confocal microscope. Endogenous AgMOBP1 displayed heterogeneous expression among individual midgut epithelial cells but was consistently enriched toward the apical region (**Fig. 2C,D,E**). Signal intensity increased from the basal plane toward the lumen-facing surface, where AgMOBP1 appeared as punctate clusters associated with the apical membrane and microvillar region. A subset of cells also showed lateral and top membrane staining as seen through a midgut cross-section cut (**Fig. 2D**). The signals did not appear in the negative control (**Supplemental Fig. S6**). Its level was quantified using intensity pixel, showing the significant localization observed on the apical side and its membrane (**Supplemental Fig. S7**). Three-dimensional visualization further revealed surface-associated clustered structures (**Fig. 2F**). Furthermore, we have visually observed a high intensity of the signal and an increased presence of AgMOBP1 clusters in regions co-located with bacterial entities markers at midgut luminal side, as indicated by DAPI staining (**Fig. 2G,H**). These data indicate that AgMOBP1 is selectively targeted to the luminal interface of midgut epithelial cells, positioning it at the site of parasite contact during invasion (**Fig. 2I**).

### AgMOBP1 directly interacts with a sexual-stage *Plasmodium* protein identified as PfSyn5

Given the localization and functional role of AgMOBP1, we hypothesized that it engages a parasite ligand during mosquito infection. To test this, recombinant AgMOBP1, produced through baculovirus expression system (**Supplemental Fig. S8-10**), was assayed for binding to sexual-stage *P. falciparum*. In ELISA, AgMOBP1 bound robustly to immobilized parasite proteins (**Fig. 3A**). Far-western blotting (**Fig. 3C**) identified a prominent interacting species of approximately 34 kDa (**Fig. 3B**), with strongest enrichment in the insoluble fraction containing membrane- and cytoskeleton-associated proteins, whereas little binding was observed in the soluble fraction.

**Figure 3.**
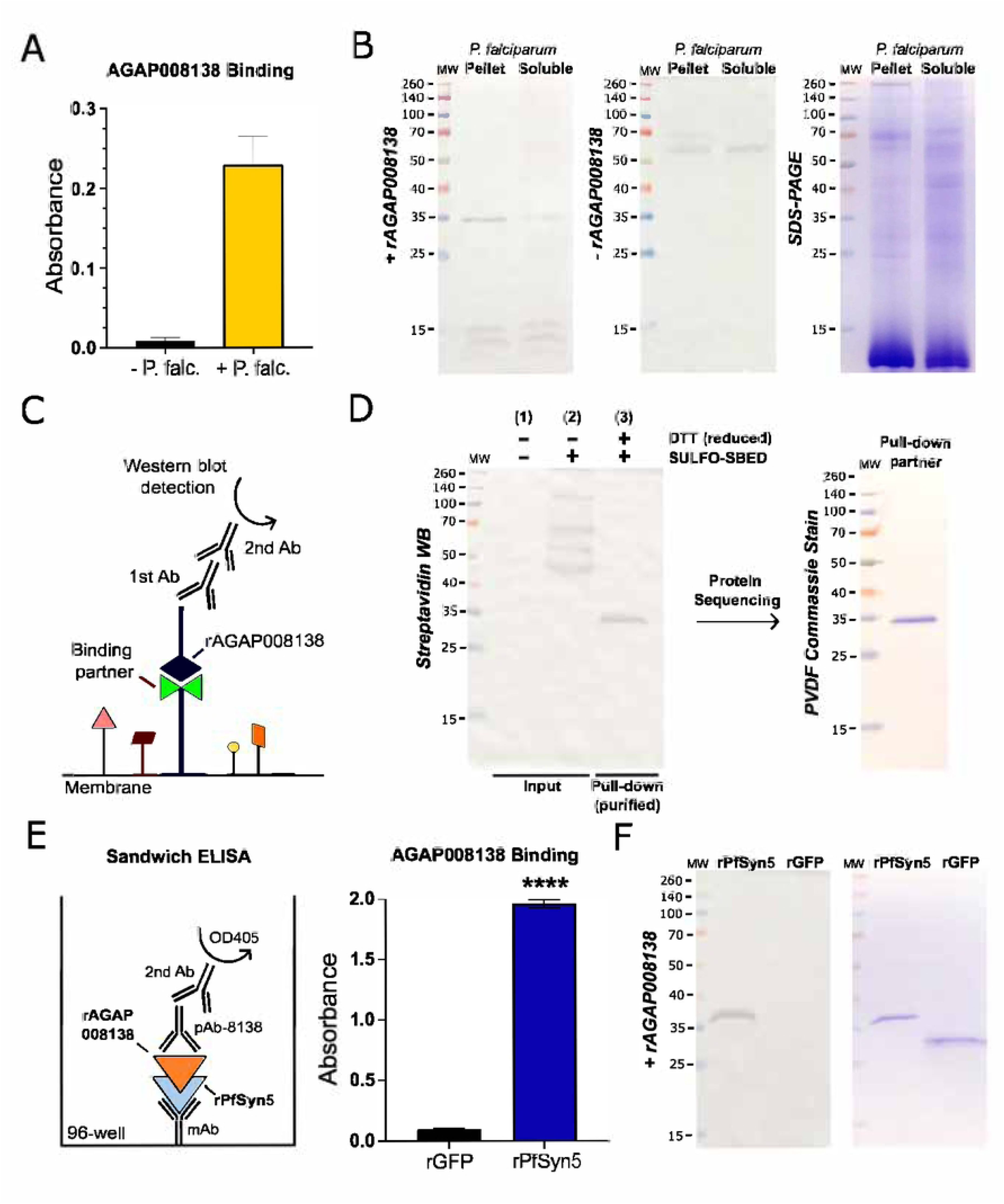
rAgMOBP1 binding to a specific protein in *P. falciparum* lysates and whole cells. **A.)** ELISA binding of rAgMOBP1 to proteins in the *P. falciparum* homogenate (P. falc.). **B.)** Far-western blot analysis showing the binding of AgMOBP1 to *P. falciparum* proteins in the membrane (pellet) and cytosol (soluble) fractions, separated by SDS-PAGE. A control blot without AgMOBP1 (-rAGAP008138) was included to validate specific binding interactions. **C.)** Far-western blotting concept illustration. The SDS-PAGE separated proteins are probed with recombinant (r) AgMOBP1 and then with anti-AgMOBP1 pAb, followed by secondary antibody and detection. **D.)** Sulfo-SBED labeled rAgMOBP1 interacting with a specific protein using the intact sexual stage culture of the *P. falciparum* parasite in a double pulldown assay. **Left**, Streptavidin western blot identifying a single protein of interest. **Right**, the single band blotted onto a PVDF membrane for Edman degradation sequencing. **E.)** Sandwich ELISA of rAGMOBP1 binding to rPfSyn5. **F.)** Far-western blot of rAgMOBP1 binding to rPfSyn5 (left), Coomassie staining of the same samples run in an SDS-PAGE (right). rGFP is a negative control, as it does not bind to rAgMOBP1.

To validate this interaction independently, we performed cross-linking pulldown assays using intact sexual-stage parasites (**Supplemental Fig. 11**) and the trifunctional reagent Sulfo-SBED. These experiments again detected a specific interacting protein migrating at ∼34 kDa (**Fig. 3D**, **Supplemental Fig. S12**), consistent with the far-western result. Subsequent sequential purification through double pulldown, band isolation, and protein identification by N-terminal sequencing revealed a sequence of “PYVDKTEEFF” (**Supplemental Dataset S3**). This sequence matches a *P. falciparum* syntaxin-5–like protein, PfSyn5 (PF3D7_1332000, XP_001350051.1, Qa-SNARE) (**Supplemental Fig. S13)**.

We next generated recombinant PfSyn5 in insect cells (**Supplemental Fig. S14-16**) and tested direct binding. AgMOBP1 bound immobilized recombinant PfSyn5 in ELISA (**Fig. 3E**) and recognized purified PfSyn5 in far-western assays, whereas negative controls did not (**Fig. 3F**). Together, these complementary biochemical approaches identify PfSyn5 as a specific parasite binding partner of AgMOBP1.

### Structural and functional features of PfSyn5

PfSyn5 is encoded by a five-exon gene on chromosome 13 and produces a 281-amino-acid protein with a predicted mass of 32.8 kDa (**Fig. 4A**). Unlike AgMOBP1, PfSyn5 lacks a signal peptide but contains a C-terminal transmembrane helix and a conserved t-SNARE domain, consistent with a membrane trafficking protein. Structural modeling predicted a predominantly α-helical architecture with multiple coiled-coil elements, typical of SNARE-family proteins (**Fig. 4B**).

**Figure 4.**
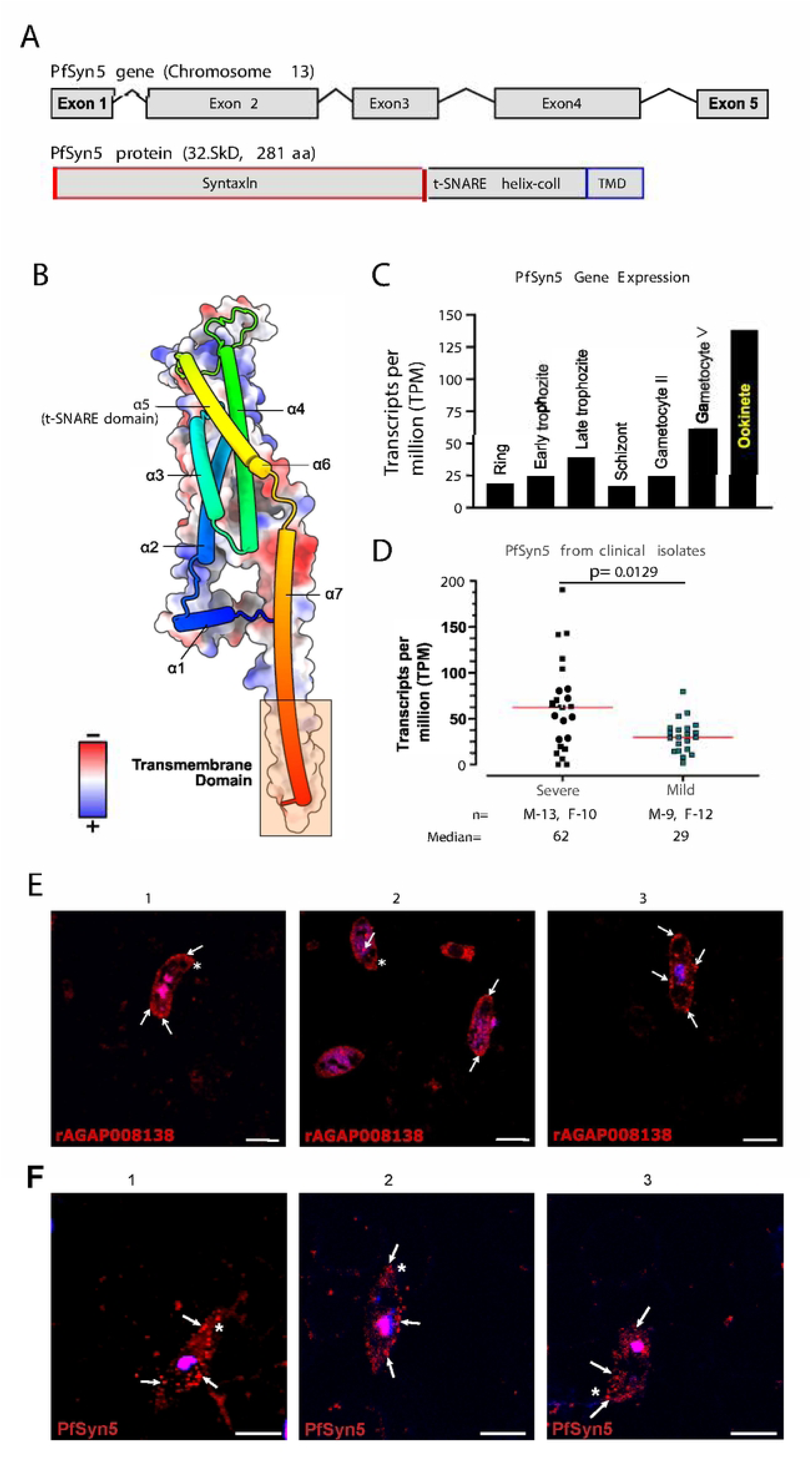
PfSyn5 characteristics, structure, and expression. **A.)** PfSyn5 has five exons coding for a 281 amino acid long protein with an estimated size of 32.8kD. PfSyn5 has three domains: the syntaxin, the t-SNARE helix-coil, and the transmembrane domain at the C-terminal. **B.)** AlphaFold2 prediction of PfSyn5 shows seven alpha-helices, where number five is the t-SNARE domain. PfSyn5 has more positively charged residues, which are essential for the recruitment and clustering of SNARE complexes, promoting membrane deformation and facilitating the fusion of vesicles with target membranes. **C.)** PfSyn5 expression profiles from different *Plasmodium* life cycle stages, adapted from a previous study. PfSyn5 is highest expressed in the ookinete stage. **D.)** A study examining clinical *Plasmodium* isolates in malaria patients, categorized by severity (severe and mild outcomes). PfSyn5 is expressed more in patients with severe malaria. **E.)** Subcellular location of PfSyn5 via indirect detection of rAgMOBP1 interaction. **F.)** Direct subcellular location detection of PfSyn5 via anti-PfSyn5 pAb. White arrows indicate distinct localizations within the ookinete, close to the membrane. Scale bars, 5µm.

Reanalysis of published functional genomics datasets indicated that PfSyn5 behaves as an essential parasite gene (*40*) with a Mutagenesis Index Score (MIS) of 0.12 and a Mutant Fitness Score (MFS) of -2.83 (**Supplemental Fig. S17**). Transcriptomic data across parasite life stages (*41*) showed expression throughout development, with marked enrichment during the ookinete stage relative to asexual stages and gametocytes. The expression of PfSyn5 in the ookinete stage was approximately 5.5-fold higher than in the asexual stages and 3.2-fold higher than in the gametocyte stages, supporting a specialized role during mosquito infection (**Fig. 4C**). Additional clinical transcriptome datasets (*42*) from 23 patients with severe malaria and 21 patients with uncomplicated (mild) malaria suggested elevated PfSyn5 expression in severe malaria isolates relative to uncomplicated cases, indicating broader biological importance beyond transmission stages (**Fig. 4D**). These findings support the crucial involvement of PfSyn5 in the pathogenesis of malaria, emphasizing its significant role not only on sexual stage transmission but also in disease progression and severity in the asexual stages.

Immunofluorescence analysis of sexual-stage parasites using indirect detection with AgMOBP1 binding revealed punctate intracellular PfSyn5 localization concentrated beneath the cytoplasmic membrane and enriched near apical regions (**Fig. 4E**). This vesicular pattern was also observed both by direct staining with anti-PfSyn5 antibodies. (**Fig. 4F**), consistent with localization to secretory or trafficking vesicles (*43*).

### Polyclonal antibodies against PfSyn5 potently blocked malaria transmission

Because AgMOBP1 promotes infection and directly binds PfSyn5, we tested whether blocking PfSyn5 interferes with parasite transmission. Purified anti-PfSyn5 polyclonal antibodies were mixed with infectious *P. falciparum* cultures and administered to mosquitoes by standard membrane feeding assay. Anti-PfSyn5 antibodies nearly completely abolished oocyst formation at concentrations of 1 µg/mL and above (**Fig. 5A**). At lower doses, inhibition remained strongly concentration dependent (**Fig. 5B**).

**Figure 5.**
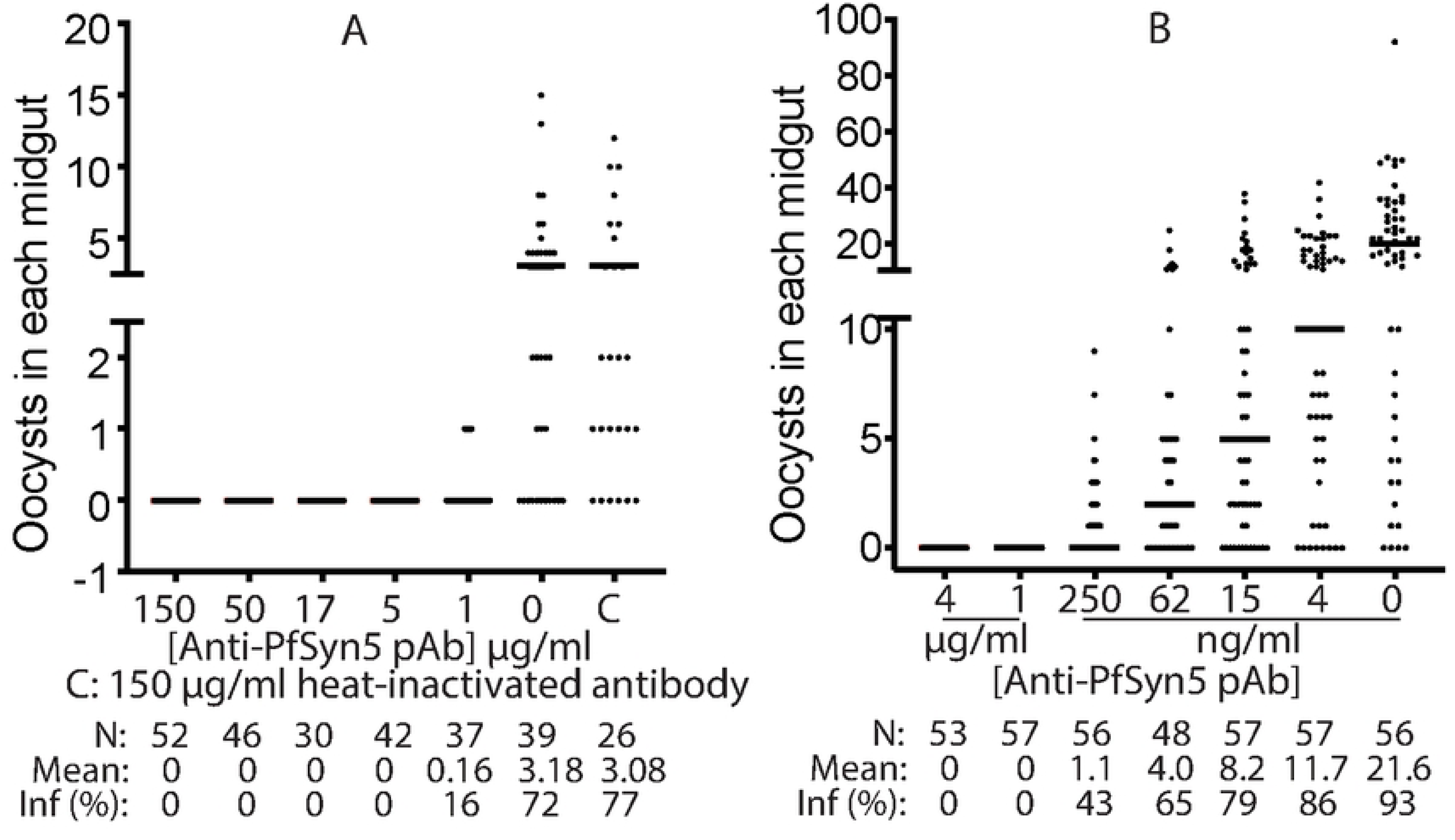
Anti-PfSyn5 polyclonal antibodies inhibited *P. falciparum* transmission to *An. gambiae* in a concentration-dependent manner. **A.**) Anti-PfSyn5 completely blocked *P. falciparum* infection in *An. gambiae* midguts at high concentrations (≥1 μg/ml). The results represent three replicates. **B.**) Different low concentrations of Anti-PfSyn5 inhibited *P. falciparum* infection in *An. gambiae* midguts in a concentration-dependent manner. IC50 of Anti-PfSyn5 is 3.2 ng/ml. Same experiments were conducted three times and show similar results.

Dose–response analysis (**Fig. 5B**) yielded an estimated IC_50_ of approximately 3.2 ng/mL (∼20 pM), indicating exceptionally high potency. This concentration corresponds to roughly one-millionth of the total IgG concentration normally present in human blood. The magnitude of inhibition exceeded that observed with anti-AgMOBP1 antibodies, suggesting that PfSyn5 is a particularly vulnerable node in the transmission process. These findings establish PfSyn5 as an unusually powerful target for transmission-blocking intervention and support the model that the AgMOBP1–PfSyn5 interaction is functionally required for successful mosquito infection.

## Discussion

This study identifies a previously uncharacterized mosquito–parasite molecular interaction that is required for efficient malaria transmission. We define AgMOBP1 as a membrane-associated midgut factor in *Anopheles gambiae* that promotes *Plasmodium falciparum* infection and identify the parasite syntaxin-5–like protein PfSyn5 as its binding partner. Functional disruption of either side of this interface reduced parasite transmission, whereas antibodies targeting PfSyn5 exhibited exceptionally potent transmission-blocking activity. Together, these findings establish the AgMOBP1–PfSyn5 axis as a previously unrecognized and actionable vulnerability in the mosquito stage of the malaria life cycle.

A notable feature of AgMOBP1 is its unconventional membrane association (*44*). Although the protein lacks a canonical transmembrane helix, it consistently partitioned with membrane fractions and localized to the apical surface of midgut epithelial cells. The presence of a CaaX motif near the C terminus suggests that post-translational prenylation may contribute to membrane anchoring (*39, 45*), although direct biochemical validation will be required. More broadly, peripheral membrane proteins can associate with lipid bilayers through lipidation, electrostatic interactions, amphipathic helices, or binding to integral membrane partners (*46–49*). Our data are most consistent with AgMOBP1 functioning as a surface-associated scaffold or receptor-like factor positioned at the luminal interface where invading ookinetes contact the epithelium.

The regulation of AgMOBP1 expression further supports a physiological role at the mosquito midgut surface. AgMOBP1 was strongly induced by blood feeding and reached peak abundance during the period in which ookinetes traverse the midgut epithelium. Expression was further elevated after infectious blood meals, suggesting that parasite presence either directly or indirectly amplifies host responses at the invasion site. One interpretation is that AgMOBP1 normally participates in post-feeding epithelial remodeling, nutrient uptake, membrane turnover, or barrier maintenance, and that parasites exploit this host program to facilitate invasion. Such co-option of host physiology is a common feature of pathogen entry strategies across biological systems (*16*).

Our localization data provide additional mechanistic context. Endogenous AgMOBP1 was enriched at the apical membrane and microvillar region, precisely where ookinetes first encounter the midgut epithelium after crossing the peritrophic matrix. This spatial positioning strongly supports a direct role in host–parasite contact rather than a distal systemic effect. The punctate and clustered distribution of AgMOBP1 may reflect organization into membrane microdomains or higher-order complexes that concentrate binding activity at discrete surface sites. Determining whether these structures correspond to lipid rafts, endocytic zones, or cytoskeletal assemblies will be an important direction for future work.

The identification of PfSyn5 as the parasite ligand is particularly important. Syntaxin-family proteins are best known as SNARE components that mediate vesicle docking and membrane fusion in eukaryotic cells (*50*). Consistent with this biology, PfSyn5 contains a predicted transmembrane segment and t-SNARE domain and localized to punctate intracellular structures concentrated near the parasite periphery and apical region. These observations are compatible with association with trafficking vesicles, secretory organelles, or Golgi-derived compartments (*51*). Elevated expression during the ookinete stage further suggests that PfSyn5 has a specialized role during mosquito infection, when directional secretion and membrane remodeling are essential for motility and invasion.

How might the AgMOBP1–PfSyn5 interaction promote transmission? Several non-mutually exclusive models are plausible. First, PfSyn5-containing vesicles or membrane domains may transiently engage AgMOBP1 at the parasite–host interface to facilitate adhesion and stable contact during epithelial traversal. Second, this interaction could promote localized membrane remodeling or fusion-like events that enhance parasite entry. Third, PfSyn5 may regulate secretion of effector cargoes required for epithelial disruption, immune evasion, or traversal, with AgMOBP1 acting as a host docking factor. Our current data support a functional interaction but do not yet distinguish among these mechanisms. High-resolution live imaging, conditional parasite genetics, and structure-guided mutagenesis will be valuable next steps.

From a translational perspective, PfSyn5 is the most striking outcome of this study. Anti-PfSyn5 polyclonal antibodies completely blocked transmission at low microgram-per-milliliter concentrations and displayed an estimated IC_50_ in the low nanogram-per-milliliter range. This level of activity compares favorably with leading transmission-blocking antibody benchmarks such as Psf48/45 (*52–54*) and indicates that PfSyn5 is highly accessible or functionally indispensable during the relevant stage of infection. Because only a fraction of total polyclonal IgG is expected to be PfSyn5-specific, the true potency of optimal monoclonal antibodies directed against vulnerable PfSyn5 epitopes may be even greater. These findings position PfSyn5 as a compelling candidate for transmission-blocking vaccines, monoclonal antibody development, or other biologic interventions.

This study has several limitations. Although multiple orthogonal assays support AgMOBP1–PfSyn5 binding, direct structural resolution of the complex is not yet available. The mechanism of AgMOBP1 membrane anchoring remains inferred rather than experimentally demonstrated. Likewise, the precise subcellular localization and essentiality of PfSyn5 during mosquito-stage development require genetic validation in parasites. Finally, transmission assays were performed under controlled laboratory conditions and should be extended to additional parasite isolates, mosquito species, and semi-field settings.

In summary, we identify AgMOBP1 as a mosquito midgut factor that facilitates malaria infection and uncover PfSyn5 as its parasite binding partner and an exceptionally potent transmission-blocking target. These findings expand the molecular framework of mosquito-stage malaria biology and reveal a new intervention axis at the vector–parasite interface. Beyond their immediate translational relevance, they underscore how unexplored mosquito and parasite proteins can expose fundamental mechanisms of transmission and create new opportunities for malaria elimination.

## Materials and Methods

### Sequence analysis and phylogenetics

*Anopheles gambiae* sequences were retrieved from VectorBase (https://vectorbase.org/vectorbase/app), and *Plasmodium falciparum* sequences were obtained from PlasmoDB (https://plasmodb.org/plasmo/app). The AgMOBP1 protein sequence was queried against OrthoDB to identify orthologs. Homologous sequences (**Supplemental Dataset S1**) were aligned using Clustal Omega and analyzed in MEGA11 (*55*). Phylogenetic relationships were inferred using the neighbor-joining method with 1,000 bootstrap replicates. Bootstrap values >40 were displayed. Domain architecture was assessed using InterPro (*56*).

### Mosquito rearing

The *Anopheles gambiae* G3 strain was obtained from BEI Resources (BeiResources.org) and maintained at 27°C, 80% relative humidity, under a 12-h light/12-h dark cycle. Larvae were fed ground koi fish food daily. Adults were maintained on 10% sucrose ad libitum. For colony propagation, females were fed human blood mixed 1:1 (v/v) with heat-inactivated human serum obtained from commercial blood suppliers.

### *P. falciparum* culture and ookinete induction

The NF54 strain of *P. falciparum* was obtained from BEI Resources and cultured in RPMI-1640 (Gibco, Amarillo, TX) complete medium supplemented with human erythrocytes (4% hematocrit), 10% AB+ serum, and 12.5 µg/mL hypoxanthine at 37°C under candle-jar conditions. Mature day-15 gametocyte cultures were used for mosquito infections or ookinete induction. For ookinete production, cultures were washed three times in complete RPMI-1640, concentrated, and resuspended in RPMI-1640 containing 20% AB+ serum, 50 µg/mL hypoxanthine, and 2 g/L NaHCO_3_. Cultures were incubated at 27°C for 24–32 h, and ookinete formation was verified by Giemsa staining.

### RNA extraction and cDNA generation

For parasite gene expression studies, ookinete cultures were processed using the PureLink RNA Mini Kit (ThermoFisher, Rockford, IL) with on-column DNase treatment. First-strand cDNA was synthesized using the RevertAid First Strand cDNA Synthesis Kit according to the manufacturer’s instructions (ThermoFisher, Rockford, IL). For mosquito samples, total RNA from dissected midguts was extracted using the PureLink RNA Mini Kit, and cDNA was generated using the Verso cDNA Synthesis Kit.

### Quantification of AgMOBP1 expression in mosquito midguts

Female mosquitoes were collected at 0, 8, 16, 24, and 32 h after blood feeding or infectious feeding. Fully engorged females were selected, and 25 midguts were dissected per replicate under ice-cold PBS. Samples were stored at −80°C until processing. Relative AgMOBP1 transcript abundance was measured by RT-PCR using the 40S ribosomal protein S7 gene as the internal reference. Reactions were performed with gene-specific primers and Phusion polymerase under standard cycling conditions. Relative expression of AgMOBP1 was assessed using the 40S S7 ribosomal protein gene as the internal reference. The primers were 8138 RT PCR-F 5‘-ATGCTGCTAAAATCGGCAC-3’,8138 RT PCR-R 5‘-CTAAACAGCCACCGGTCC-3’, S7 40S RT PCR-F 5‘-ATGGTGTTCGGTTCCAA GG-3’ and S7 40S RT PCR-R 5’-TTACAGGTAGTTCTCTGGGAATTC-3’. Three biological replicates were analyzed for each condition. PCR was performed under standard conditions using the Phusion plus DNA polymerase (ThermoFisher, Rockford, IL). 0.5µM of each primer was used with an initial denaturation step of 1 min at 98°C and then 45 cycles of 10 s at 98°C, 10 s at 60°C, and 45 s at 72°C, with a final extension of 5 min at 72°C.

### Midgut protein extraction and fractionation

Mosquito midguts were collected from sugar-fed or blood-fed females at the indicated time points (0 h-native, 8 h, 16 h, 24 h, and 32 h) and homogenized in solubilization buffer (15 mM Tris-HCl, pH 8.0, 150 mM NaCl, 5 mM EDTA, protease inhibitors, 2 µL per midgut). Samples were first centrifuged at 500 × g to remove debris. Supernatants were then centrifuged at 20,000 × g for 1 h at 4°C to separate soluble and membrane-associated fractions. The membrane pellet was resuspended in solubilization buffer containing 0.5% Triton X-100 and DDM. Fractions were analyzed by SDS-PAGE and western blotting.

### Recombinant protein expression and purification

Unless otherwise indicated, recombinant proteins were expressed in insect cells using baculovirus expression system with a C-terminal 6xHis tag. Codon-optimized AgMOBP1 and PfSyn5 constructs (**Supplemental Dataset S2**) were cloned into pFastBac vectors using BamHI and SphI as site-specific restriction enzymes (**Supplemental Table S1**). According to previous literature and to increase protein expression in insect cells (*57*), the L21 region of “AACTCCTAAAAAACCGCCACC” was added before the “ATG” of each gene. Recombinant bacmids were generated in DH10Bac cells and used to produce baculovirus stocks. High-titer virus was amplified in *Spodoptera frugiperda 9* (Sf9) cells and used to infect High Five (Hi5) cells. AgMOBP1 was expressed at a multiplicity of infection (MOI) of 10, and PfSyn5 at an MOI of 5. Cells were harvested 80 h after infection and lysed in buffer containing 500mM NaCl, 50mM sodium phosphate, 50mM imidazole, 10% glycerol, 1% Triton-X100, 1% DDM, 0.5% Tween-20, and protease inhibitors. Soluble lysates were purified by nickel affinity chromatography on HisTrap HP columns using an ÄKTA Pure system. Purified proteins were assessed by SDS-PAGE and stored at −80°C.

### Antibody production and purification

Recombinant AgMOBP1 and PfSyn5 proteins were used to generate rabbit polyclonal antibodies (BosterBio, Pleasanton, CA). Immunization schedules included primary immunization with Freund’s complete adjuvant followed by booster injections with Freund’s incomplete adjuvant. Serum IgG was purified using protein A-affinity chromatography. Sodium azide was removed by centrifugal filtration before functional assays.

### Standard membrane feeding assays

Transmission-blocking assays were performed using standard membrane feeding assays (SMFAs). Purified antibodies or recombinant proteins were exchanged into PBS, quantified by A280 and BCA assay, and added to infectious blood meals containing mature *P. falciparum* gametocytes, erythrocytes, and heat-inactivated human serum. Samples were fed to mosquitoes through glass membrane feeders maintained at 37°C. Fully engorged females were separated and maintained on 10% sucrose. Midguts were dissected 7 d post-feeding, stained with 0.1% mercurochrome, and oocysts were counted microscopically.

### Immunofluorescence assays of mosquito midguts

Mosquito midguts were dissected in ice-cold PBS 24 h after feeding and carefully opened to remove the blood bolus. Tissues were fixed in 4% paraformaldehyde, permeabilized in PBS containing Triton X-100, and blocked in PBS containing BSA and gelatin. Samples were incubated overnight at 4°C with anti-AgMOBP1 primary antibody, followed by Alexa Fluor 594–conjugated secondary antibody. Slides were mounted in antifade medium containing DAPI and imaged using an Olympus Fluoview confocal microscope. Images were processed using Fiji/ImageJ and 3D Viewer.

### Immunofluorescence assay of sexual-stage parasites

Ookinete cultures were deposited onto coverslips and fixed in 4% paraformaldehyde in PBS at room temperature for 30 min. Cells were quenched with 100 mM glycine for 20 min, blocked in 2% BSA in PBS for 90 min, and incubated with anti-PfSyn5 (5 μg/mL in PBS) antibody or recombinant AgMOBP1 (10 μg/mL in PBS) followed by anti-AgMOBP1 antibody. Alexa Fluor 594–conjugated secondary antibodies (2 μg/mL in PBS; Invitrogen) were used for detection. Coverslips were mounted and imaged by confocal microscopy with sequential channel acquisition.

### Western blotting

Proteins were separated by SDS-PAGE and transferred to nitrocellulose membranes. Membranes were blocked in milk-containing buffer and incubated with primary antibodies overnight at 4°C. After washing, membranes were incubated with alkaline phosphatase–conjugated secondary antibodies. Signals were developed using NBT/BCIP substrate. Streptavidin was used where indicated to detect biotin-labeled proteins.

### ELISA-based binding assays

For parasite lysate binding assays, 96-well plates were coated with sexual-stage parasite lysates (1mg/ml proteins, 50µL/well) overnight at 4°C. Plates were blocked with BSA and incubated with recombinant AgMOBP1. Bound protein was detected using anti-AgMOBP1 antibodies and alkaline phosphatase–conjugated secondary antibodies. For recombinant protein interaction assays, anti-His antibody (1 µg/mL) was first immobilized, followed by capture of recombinant PfSyn5 and incubation with recombinant AgMOBP1. Absorbance was measured at 405 nm.

### Far-western blotting and cross-linking assays

Percoll-enriched (35/65% Percoll gradient) sexual-stage parasites (*58*) were lysed ((10 mM Tris-HCl, pH 7.4, 150 mM NaCl, 1 mM EDTA, protease Inhibitor) and separated into soluble and insoluble fractions. Proteins were resolved by SDS-PAGE and transferred to nitrocellulose membranes. After denaturation and stepwise guanidine-HCl refolding (*59*), membranes incubated in blocking buffer (0.1% Tween-20, 1X PBS, and 5% skim milk containing 0.1 mM levamisole). The membranes were incubated with 1 µM recombinant AgMOBP1 as bait protein (100 mM NaCl, 20 mM Tris (pH 7.6), 0.5 mM EDTA, 10% glycerol, 0.1% Tween-20, 2% skim milk powder). Protein interactions were stabilized by EDC cross-linking, and bound AgMOBP1 was detected immunologically. Interacting bands were excised for downstream protein identification.

### Sulfo-SBED labeling and pulldown identification of binding partners

AgMOBP1-expressing insect cells were labeled with the membrane-impermeable trifunctional cross-linker Sulfo-SBED. Labeled cells or purified recombinant AgMOBP1 were incubated with intact or lysed parasites, followed by UV activation to transfer biotin to interacting proteins. Samples were purified sequentially by Ni-NTA affinity and streptavidin pulldown. Enriched proteins were analyzed by SDS-PAGE, western blotting, mass spectrometry, or Edman degradation sequencing.

## Availability of data and material

All data generated or analyzed during this study are included in the manuscript, Supporting Information, and Datasets.

## Author contributions

**JR:** Designed experiments, Executed experiments, Data collection, Data analysis, Data interpretation, Drafted manuscript. **GN:** Parasite culture, initial training and conducting experiments. **LK:** Assisted with experimental execution. **LL**: Conducting protein expression experiments. **XW:** Bioinformatics and insectary support. **JL:** Conceptualization, Experimental Design, Execution, and Data Interpretation, Editing, and Supervision.

## Acknowledgments

We want to acknowledge the TA and fellowship support from the Biomedical Sciences Ph.D. program, Drs. Hackett and Francesco Zarate from the Dr. John Hackett lab for their occasional support in troubleshooting, Dr. Molina-Cruz from the Laboratory of Malaria and Vector Research at the National Institutes of Health (NIH) for all the practical advice and external consulting, and Drs. Kevin Chandler and Carlos Pavan help with nLC-MS/MS experiments. Dr. Prem Chapagain helps on protein structure prediction by AlphaFold. Many graduate students and staff provided various forms of assistance during Julian Ramelow’s graduate studies.

